# Identifying potential hosts of short-branch Microsporidia

**DOI:** 10.1101/2020.08.13.249938

**Authors:** Annemie Doliwa, Micah Dunthorn, Erika Rassoshanska, Frédéric Mahé, David Bass, Camila Duarte Ritter

## Abstract

Microsporidia are obligate parasites that are closely related to Fungi. While the widely-known “long-branch” Microsporidia infect mostly animals, the hosts of “short-branch” Microsporidia are only partially characterized or not known at all. Here, we used network analyses from Neotropical rainforest soil metabarcoding data, to infer co-occurrences between environmental lineages of short-branch microsporidians and their potential hosts. We found significant co-occurrences with several taxa, especially with Apicomplexa, Cercozoa, Fungi, as well as some Metazoa. Our results are the first step to identify potential hosts of the environmental lineages of short-branch microsporidians, which can be targeted in future molecular and microscopic studies.

---

Environmental DNA sequencing studies have uncovered numerous protistan parasitic groups in different environments. For example, apicomplexans can dominate soils in Neotropical rainforests (Mahé et al., 2017) and Syndiniales can likewise be species-rich in marine waters (de Vargas et al., 2015). At least at larger taxonomic levels, it is relatively straightforward to infer the hosts of these protistan parasites: the apicomplexans mostly infect animals (Votýpka et al., 2017) and the Syndiniales infect metazoans and other protists (Guillou et al., 2008). However, we do not always know so clearly who are the hosts for other protistan parasite groups uncovered in environmental DNA sequencing studies. One such example of this lack of knowing who are the potential hosts are the “short-branch” microsporidians (Bass et al., 2018).

The short-branch microsporidians form a basal grade leading up to the more widely-known “long-branch” microsporidia (Bass et al., 2018). Long-branch microsporidia are mostly parasites of metazoans (Cali et al., 2017), but some can infect ciliates and other protists (Fokin et al., 2008). While the long-branch microsporidia have highly reduced genomes and complex polar filaments that allow to penetrate cells, the short-branch microsporidians have less reduced genomes and they lack fully-developed polar filaments (Bass et al., 2018). The short-branch microsporidians include the partially-characterized *Paramicrosporidium* that are parasites of *Saccamoeba limax* (Michel et al., 2009) and *Vannella* (Michel et al., 2000), *Mitosporidium* that are parasites of the crustacean Daphnia (Haag et al., 2014), as well as *Morellospora, Chytridiopsis*, and the metchnikovellids. The short-branch microsporidians also include numerous environmental lineages recently uncovered in a re-analysis of a metabarcoding study of Neotropical rainforest soils (Bass et al., 2018). Presumably all of these environmental lineages phylogenetically assigned to the short-branch microsporidians are likewise parasitic; it is unknown, though, who are their potential microbial- or macro-organismic hosts, or where to even begin to look for them in environments as species-rich as tropical forests.

A novel approach to evaluating the diversity of protistan parasites and their hosts in metabarcoding datasets was recently demonstrated (Singer et al., 2020). Using linear regression models, Singer et al. (2020) showed that the abundances of apicomplexans and their metazoan hosts positively correlated across alpine sites in Switzerland. That type of analysis is dependent in part, though, on knowing what are the potential hosts through previous observations. Another approach to unravel potential host-parasite relationships when the hosts are unknown, is to use co-occurrence network analyses. Although network analyses based on co-occurrences do not confirm biotic interactions (Blanchet et al. 2020), cooccurrence network can highlight potentially interesting taxonomic groups as potential hosts.

We used a co-occurrence network built from metabarcoding data from Mahé et al. (2017). Briefly, the data came from soils collected in lowland rainforests in Costa Rica, Panama, and Ecuador. The soils were amplified using broad eukaryotic primers for the V4 region of SSU-rRNA (Stoeck et al., 2010) and sequenced with Illumina MiSeq. After initial cleaning steps, reads were clustered into operational taxonomic units (OTUs) with Swarm (Mahé et al., 2015) and taxonomically assigned using the PR^2^ database (Guillou et al., 2013). Most of the OTUs were assigned to different protistan taxa, while others were assigned to the Fungi and Metazoa. From this original data, refinements of the taxonomic assignments placed 974 OTUs into the short-branch microsporidia (Bass et al., 2018). We calculated the richness of these OTUs, and compared the shared OTUs by country using vegan v.2.5-6 (Oksanen, 2011) in R v.3.6.3 (R Core Team, 2020).

Representative sequences from all eukaryotic OTUs were used to construct a cooccurrence network with the NetworkNullHPC script (https://github.com/lentendu/NetworkNullHPC). In this network, the OTUs are represented as nodes, and a statistically significant Spearman correlation between two OTUs are represented by an edge between them. The network contains only OTUs with a significant co-occurrence with at least one other OTU, using a null model following Connor et al. (2017). The resulting classification matrices were combined in R with tidyverse v.1.3.0 (Wickham et al., 2019) and igraph v.1.2.4.2 (Csárdi and Nepusz, 2006), then explored and visualized with Gephi v.0.9.2 (Bastian et al., 2009) using the Yifan Hu layout. The network was filtered for short-branch microsporidians and their correlating nodes, then further explored with a Sankey diagram made in networkD3 v.0.4 (Allaire et al., 2017). Spearman correlation was used to link short-branch microsporidia and their correlating nodes in the Sankey diagram as it is a ponderation between the number of edges and the strength of the correlation among nodes.

The co-occurrence network consisted of 14,329 edges involving 368 nodes (approximately 2.40% of all OTUs in the dataset). Costa Rica had the highest richness of short-branch microsporidia, and the highest number of exclusive OTUs (Fig. S1 & S2). However, just 15 widespread microsporidian OTUs were present in the network, corresponding to approximately 1.54% of all their OTUs in the dataset (Fig. 1). Of these OTUs, 11 had a closest taxonomic assignment to the *Paramicrosporidium*, and four to the *Mitosporidium*, although the OTUs likely form independent environmental lineages (Table S1). Filtering the co-occurrences for correlations only associated with these 15 short-branch microsporidian OTUs resulted in 1,223 edges involving 244 nodes, with 768 edges belonging to OTUs assigned to the ‘paramicrosporidium’ and 455 edges to OTUs assigned to the ‘mitosporidium’ (Fig. 2; Tables S2 and S3). The three most prominent groups co-occurring with the short-branch microsporidians are the Cercozoa, Fungi, and Apicomplexa. The two largest groups in the cercozoans to form co-occurrences were the largely bacterivorous testate amoebae in the Thecofilosea and Euglyphida. Within the fungi, the largest groups were the Chytridiomycota and the Ascomycota, that are mostly found in those tropical soils in yeast-forming stages (Dunthorn et al., 2017). Most of the apicomplexans were in the Gregarinasina, which are parasites of invertebrates and dominated the soil protistan communities in the tropical forests (Mahé et al., 2017). Some other groups that also co-occurred with the short-branch microsporidians included: Amoebozoa, Endomyxa, Ciliophora, Metazoa, and Oomycota. The few metazoans in the networks were assigned to the Nematoda and Annelida (Table S4).

**Figure 1.**
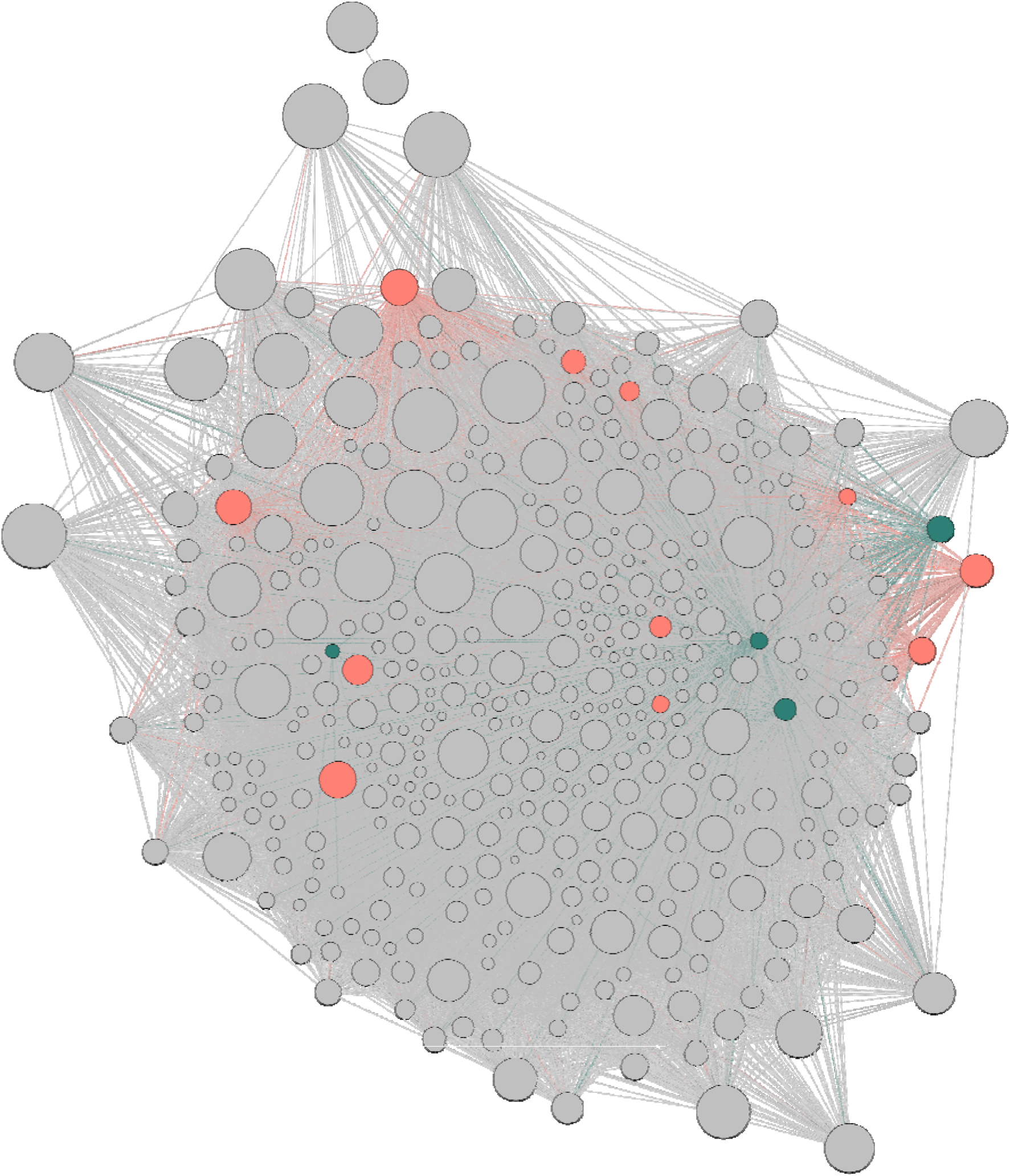
Co-occurrence network with OTUs as nodes and correlations as edges; the node size illustrates the abundance of the OTU. Mitosporidian OTUs are highlighted in turquois and Paramicrosporidian OTUs in red.

**Figure 2.**
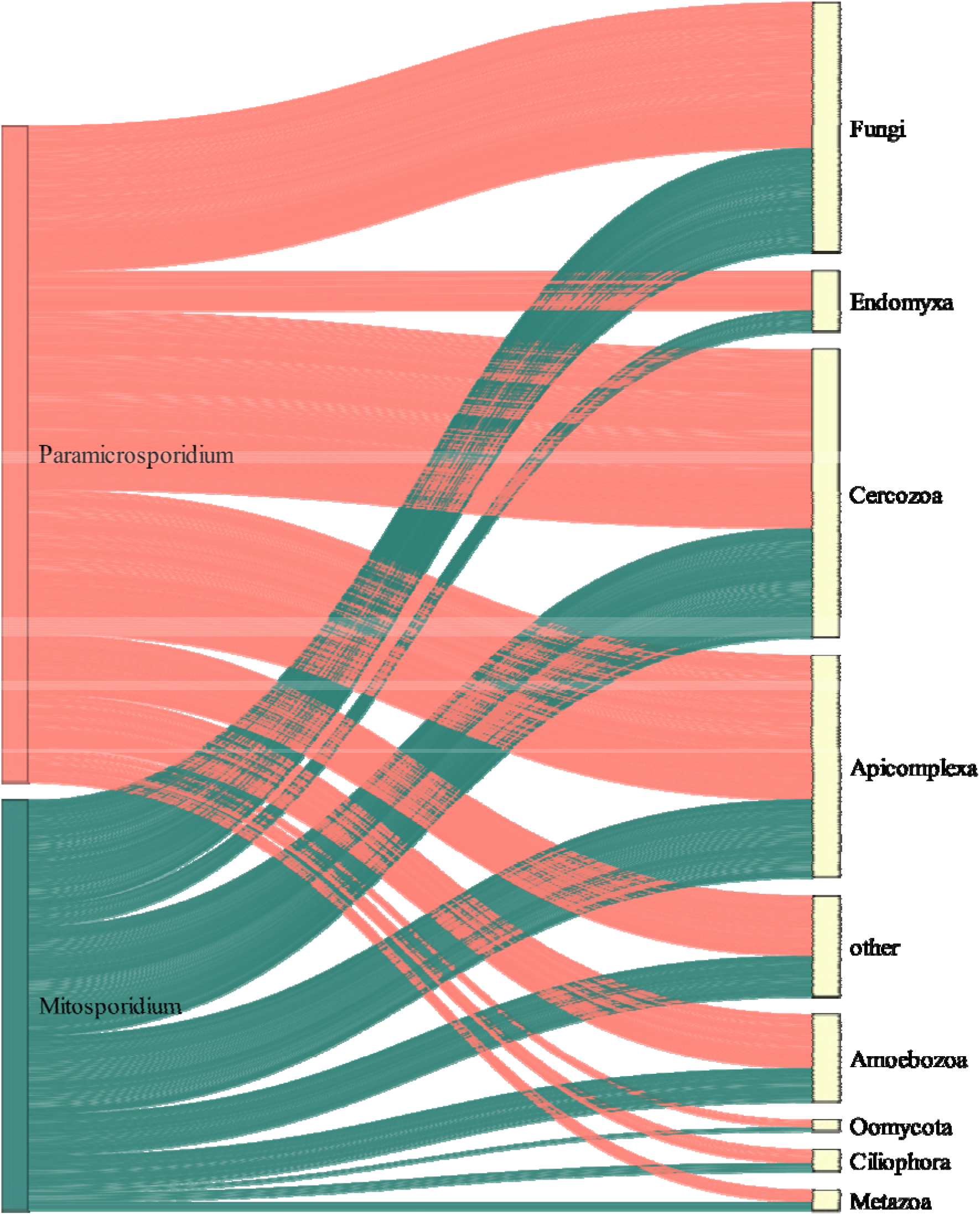
Sankey diagram showing the edges between microsporidian OTUs (left) and their target OTUs (right) in the co-occurrence network. Edges with Paramicrosporidian OTUs are marked in red, those with Mitosporidian OTUs are colored as turquoise.

Although we found protists, fungi, and metazoans co-occurring with the short-branch microsporidians, the network analyses do not directly demonstrate that they are actual hosts. Co-occurrences can be inferred because of similar environment preferences, and actual biotic interactions may not have been inferred because the signal was too weak in the data (Blanchet et al., 2020). Additionally, some of the co-occurrences here could have been inferred just because the cercozoans, fungi, and apicomplexans were extremely OTU-rich in the dataset. Potentially more of the short-branch microsporidians could have metazoan hosts, but the use of the SSU-rRNA environmental sequences likely underestimated their diversity. Yet, we inferred co-occurrences between four Annelida and one Nematode, a pattern found before with other Microsporidia (e.g., Troemel et al. 2008, Oumouna et al. 2000).

Even in light of these potential limitations, the co-occurrence networks here highlight taxa that should be evaluated further for being the hosts of the environmental linages of shortbranch microsporidians in complex Neotropical rainforest communities. These additional observations could include fluorescence *in situ* hybridization (FISH) probes designed for the short-branch microsporidians and used on cell isolates of cercozoans, fungi, apicomplexans, and possibly metazoan. Furthermore, the co-occurrence network proposed here can be used to evaluate other protists uncovered in environmental DNA sequencing studies potential hosts in other complex environments.

## Acknowledgements

Funding came from the Alexander von Humboldt Foundation to C.D.R., and the Deutsche Forschungsgemeinschaft (#DU1319/5-1) to M.D.

## References

Allaire, J.J., Gandrud, C., Russell, K., Yetman, C., 2017. networkD3: D3 Javascript Network Graphs from R.

Bass, D., Czech, L., Williams, B.A.P., Berney, C., Dunthorn, M., Mahé, F., Torruella, G., Stentiford, G.D., Williams, T.A., 2018. Clarifying the relationships between Microsporidia and Cryptomycota. Journal of Eukaryotic Microbiology 65, 773–782.

Bastian, M., Heymann, S., Jacomy, M., 2009. Gephi: An open source software for exploring and manipulating networks.

Blanchet, F.G., Cazelles, K., Gravel, D., 2020. Co occurrence is not evidence of ecological interactions. Ecology Letters 23, 1050–1063.

Cali, A., Becnel, J.J., Takvorian, P.M., 2017. Microsporidia, in: Archibald, J.M., Simpson, A.G.B., Slamovits, C.H. (Eds.), Handbook of the Protists, Second Edition. Springer International Publishing, Cham, pp. 1559–1618.

Connor, N., Barberán, A., Clauset, A., 2017. Using null models to infer microbial cooccurrence networks. PLOS ONE 12, e0176751.

Csárdi, G., Nepusz, T., 2006. The igraph software package for complex network research. InterJournal Complex Systems, 1695.

de Vargas, C., Audic, S., Henry, N., Decelle, J., Mahe, F., Logares, R., Lara, E., Berney, C., Le Bescot, N., Probert, I., Carmichael, M., Poulain, J., Romac, S., Colin, S., Aury, J.-M., Bittner, L., Chaffron, S., Dunthorn, M., Engelen, S., Flegontova, O., Guidi, L., Horak, A., Jaillon, O., Lima-Mendez, G., Luke, J., Malviya, S., Morard, R., Mulot, M., Scalco, E., Siano, R., Vincent, F., Zingone, A., Dimier, C., Picheral, M., Searson, S., Kandels-Lewis, S., Tara Oceans Coordinators, Acinas, S.G., Bork, P., Bowler, C., Gorsky, G., Grimsley, N., Hingamp, P., Iudicone, D., Not, F., Ogata, H., Pesant, S., Raes, J., Sieracki, M.E., Speich, S., Stemmann, L., Sunagawa, S., Weissenbach, J., Wincker, P., Karsenti, E., Boss, E., Follows, M., Karp-Boss, L., Krzic, U., Reynaud, E.G., Sardet, C., Sullivan, M.B., Velayoudon, D., 2015. Eukaryotic plankton diversity in the sunlit ocean. Science 348, 1261605.

Dunthorn, M., Kauserud, H., Bass, D., Mayor, J., Mahé, F., 2017. Yeasts dominate soil fungal communities in three lowland Neotropical rainforests. Environmental Microbiology Reports 9, 668–675.

Fokin, S.I., Di Giuseppe, G., Erra, F., Dini, F., 2008. *Euplotespora binucleata* n. gen., n. sp. (Protozoa: Microsporidia), a Parasite Infecting the Hypotrichous Ciliate *Euplotes woodruffi*, with Observations on Microsporidian Infections in Ciliophora. Journal of Eukaryotic Microbiology 55, 214–228.

Guillou, L., Bachar, D., Audic, S., Bass, D., Berney, C., Bittner, L., Boutte, C., Burgaud, G., de Vargas, C., Decelle, J., del Campo, J., Dolan, J.R., Dunthorn, M., Edvardsen, B., Holzmann, M., Kooistra, W.H.C.F., Lara, E., Le Bescot, N., Logares, R., Mahé, F., Massana, R., Montresor, M., Morard, R., Not, F., Pawlowski, J., Probert, I., Sauvadet, A.-L., Siano, R., Stoeck, T., Vaulot, D., Zimmermann, P., Christen, R., 2013. The Protist Ribosomal Reference database (PR2): a catalog of unicellular eukaryote Small Sub-Unit rRNA sequences with curated taxonomy. Nucleic Acids Research 41, D597–D604.

Guillou, L., Viprey, M., Chambouvet, A., Welsh, R.M., Kirkham, A.R., Massana, R., Scanlan, D.J., Worden, A.Z., 2008. Widespread occurrence and genetic diversity of marine parasitoids belonging to *Syndiniales (Alveolata)*. Environmental Microbiology 10, 3349–3365.

Haag, K.L., James, T.Y., Pombert, J.-F., Larsson, R., Schaer, T.M.M., Refardt, D., Ebert, D., 2014. Evolution of a morphological novelty occurred before genome compaction in a lineage of extreme parasites. Proceedings of the National Academy of Sciences 111, 15480–15485.

Mahé, F., de Vargas, C., Bass, D., Czech, L., Stamatakis, A., Lara, E., Singer, D., Mayor, J., Bunge, J., Sernaker, S., Siemensmeyer, T., Trautmann, I., Romac, S., Berney, C., Kozlov, A., Mitchell, E.A.D., Seppey, C.V.W., Egge, E., Lentendu, G., Wirth, R., Trueba, G., Dunthorn, M., 2017. Parasites dominate hyperdiverse soil protist communities in Neotropical rainforests. Nature Ecology & Evolution 1, 0091.

Mahé, F., Rognes, T., Quince, C., de Vargas, C., Dunthorn, M., 2015. Swarm v2: highly-scalable and high-resolution amplicon clustering. PeerJ 3, e1420.

Michel, R., Müller, K.-D., Hauröder, B., 2009. A novel microsporidian endoparasite replicating within the nucleus of Saccamoeba limax isolated from a pond. Endocytobiosis Cell Res. 19, 120–126.

Michel, R., Schmid, E.N., Böker, T., Hager, D.G., Müller, K.-D., Hoffmann, R., Seitz, H.M., 2000. Vannella sp. harboring Microsporidia-like organisms isolated from the contact lens and inflamed eye of a female keratitis patient. Parasitology Research 86, 514–520.

Oksanen, J., 2011. Multivariate analysis of ecological communities in R: vegan tutorial. R Package Version 1, 1–43.

Oliveros, J.C., 2007. VENNY. An interactive tool for comparing lists with Venn diagrams. URL https://bioinfogp.cnb.csic.es/tools/venny/index.html

Oumouna, M., El-Matbouli, M., Hoffmann, R.W., Bouix, G., 2000. Electron microscopic study of a new microsporean *Microsporidium epithelialis* sp. n. infecting *Tubifex* sp. (Oligochaeta). Folia Parasitologica 47, 257–265.

R Core Team, 2020. R: A language and environment for statistical computing. R Foundation for Statistical computing, Vienna, Austria.

Singer, D., Duckert, C., Heděnec, P., Lara, E., Hiltbrunner, E., Mitchell, E.A.D., 2020. High-throughput sequencing of litter and moss eDNA reveals a positive correlation between the diversity of Apicomplexa and their invertebrate hosts across alpine habitats. Soil Biology and Biochemistry 147, 107837.

Stoeck, T., Bass, D., Nebel, M., Christen, R., Jones, M.D.M., Breiner, H.-W., Richards, T.A., 2010. Multiple marker parallel tag environmental DNA sequencing reveals a highly complex eukaryotic community in marine anoxic water. Molecular Ecology 19, 21–31.

Troemel, E.R., Félix, M.-A., Whiteman, N.K., Barrière, A., Ausubel, F.M., 2008. Microsporidia are natural intracellular parasites of the mematode *Caenorhabditis elegans*. PLoS Biology 6, e309.

Votýpka, J., Modrý, D., Oborník, M., Šlapeta, J., Lukeš, J., 2017. Apicomplexa, in: Archibald, J.M., Simpson, A.G.B., Slamovits, C.H. (Eds.), Handbook of the Protists, Second Edition. Springer International Publishing, Cham, pp. 567–624.

Wickham, H., Averick, M., Bryan, J., Chang, W., McGowan, L., François, R., Grolemund, G., Hayes, A., Henry, L., Hester, J., Kuhn, M., Pedersen, T., Miller, E., Bache, S., Müller, K., Ooms, J., Robinson, D., Seidel, D., Spinu, V., Takahashi, K., Vaughan, D., Wilke, C., Woo, K., Yutani, H., 2019. Welcome to the Tidyverse. Journal of Open Source Software 4, 1686.

