## Supplementary material for "Identifying potential hosts of short-branch Microsporidia": Tables S1-S4 and Figures S1-S2

**Material Supplementary for:**

**Table S1.** Additional taxonomic information for the 15 SB-microsporidian OTUs in the analyzed co-occurrence network.

| | SB-microsporidian OTUs | number of OTUs |
|---|---|---|
| a | **Mitosporidium, clone=LKM15** | 4 |
| b | **Paramicrosporidium, clone=Amb_18S_1526** | 3 |
| c | **Paramicrosporidium, clone=P34.42** | 2 |
| d | **Paramicrosporidium, clone=PFB7SP2005** | 1 |
| e | **Paramicrosporidium, clone=RSC-CHU-59** | 4 |
| f | **Paramicrosporidium+saccamoebae, strain=KSL3** | 1 |

**Table S2.** Overview of the number of edges between the Mitosporidian or Paramicrosporidian and the different taxa, divided according to the individual SB-microsporidian OTUs. The letters a – f refer to the taxonomic information for the SB-microsporidian OTUs given in Table S1.

| | OTU | Oomycota | Ciliophora | Metazoa | Endomyxa | Amoebozoa | other | Apicomplexa | Fungi | Cercozoa |
|---|---|---|---|---|---|---|---|---|---|---|
| **Mitosporidium** | **a** | 1 | 2 | 1 | 4 | 6 | 4 | 14 | 16 | 15 |
| | **a** | 1 | 3 | 4 | 8 | 15 | 19 | 29 | 45 | 48 |
| | **a** | 2 | 3 | 4 | 10 | 16 | 23 | 36 | 49 | 51 |
| | **a** | 0 | 1 | 1 | 1 | 1 | 1 | 7 | 6 | 8 |
| **Paramicrosporidium** | **e** | 1 | 3 | 3 | 8 | 14 | 23 | 33 | 44 | 52 |
| | **e** | 1 | 3 | 4 | 9 | 12 | 19 | 24 | 33 | 42 |
| | **b** | 1 | 3 | 3 | 9 | 13 | 20 | 37 | 33 | 41 |
| | **e** | 1 | 2 | 1 | 4 | 6 | 4 | 14 | 16 | 17 |
| | **e** | 0 | 0 | 1 | 0 | 0 | 0 | 4 | 4 | 0 |
| | **f** | 1 | 0 | 0 | 4 | 4 | 2 | 6 | 11 | 9 |
| | **c** | 1 | 0 | 1 | 4 | 4 | 0 | 15 | 5 | 16 |
| | **d** | 1 | 2 | 0 | 4 | 4 | 0 | 15 | 11 | 11 |
| | **b** | 0 | 0 | 0 | 0 | 1 | 0 | 5 | 3 | 3 |
| | **b** | 1 | 3 | 1 | 5 | 6 | 6 | 15 | 10 | 15 |
| | **c** | 0 | 0 | 0 | 0 | 0 | 0 | 0 | 0 | 1 |

**Table S3.** Overview of the total number of edges per taxon.

| Taxon | Total number of edges | Edges with Paramicrosporidium | Edges with Mitosporidium |
|---|---|---|---|
| **Oomycota** | 12 | 8 | 4 |
| **Metazoa** | 24 | 14 | 10 |
| **Ciliophora** | 25 | 16 | 9 |
| **Endomyxa** | 70 | 47 | 23 |
| **Amoebozoa** | 102 | 64 | 38 |
| **Other** | 121 | 74 | 47 |
| **Apicomplexa** | 254 | 168 | 86 |
| **Fungi** | 286 | 170 | 116 |
| **Cercozoa** | 329 | 207 | 122 |
| **Total number** | 1,223 | 768 | 455 |

**Table S4.** Further taxonomic information about the metazoan OTUs that showed a significant correlation with the SB-microsporidian OTUs.

| Taxonomy of metazoan OTUs | Edges |
|---|---|
| Nematozoa\|Nematoda\|Enoplia\|Triplonchida\|tobrilines\|Prismatolaimidae\|g:Prismatolaimus\|* | 3 |
| Annelida\|Pleistoannelida\|Sedentaria\|core-sedentarids\|Clitellata\|*\|*\|* | 4 |
| Annelida\|Pleistoannelida\|Sedentaria\|core-sedentarids\|Clitellata\|Lumbricina\|Glossoscolecidae\|g:Pontoscolex\|Pontoscolex+spiralis | 11 |
| Annelida\|Pleistoannelida\|Sedentaria\|core-sedentarids\|Clitellata\|Enchytraeidae\|g:Hemienchytraeus\|isolate=PDW-20 | 1 |
| Annelida\|Pleistoannelida\|Sedentaria\|core-sedentarids\|Clitellata\|Enchytraeidae\|g:Hemienchytraeus\|isolate=PDW-20 | 5 |

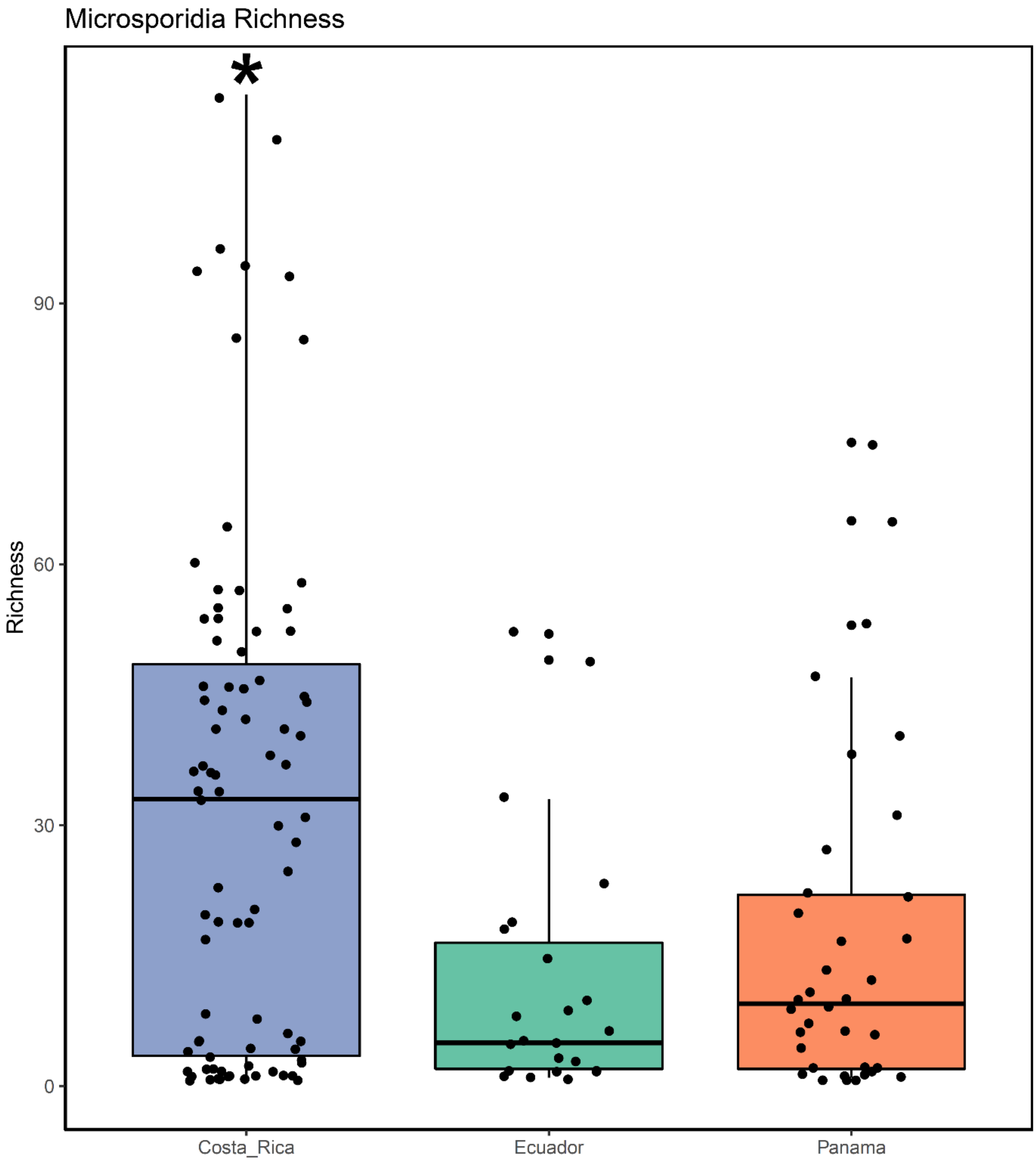

**Figure S1.** Richness of Microsporidia by country. Costa Rica has the highest and significant richness (Anova test: F value = 8.619. p = 0.0002). Turkey posthoc test to compare among countries showed significant difference between Costa Rica and Ecuador (Diff = 15.7, p = 0.005) and Costa Rica and Panama (Diff = -20.25, p = 0.002), but not between Ecuador and Panama (Diff = -4.5, p = 0.78). All analyses were calculated with vegan v.2.5-6 (Oksanen, 2011) in R v.3.6.3 (R Core Team, 2020).

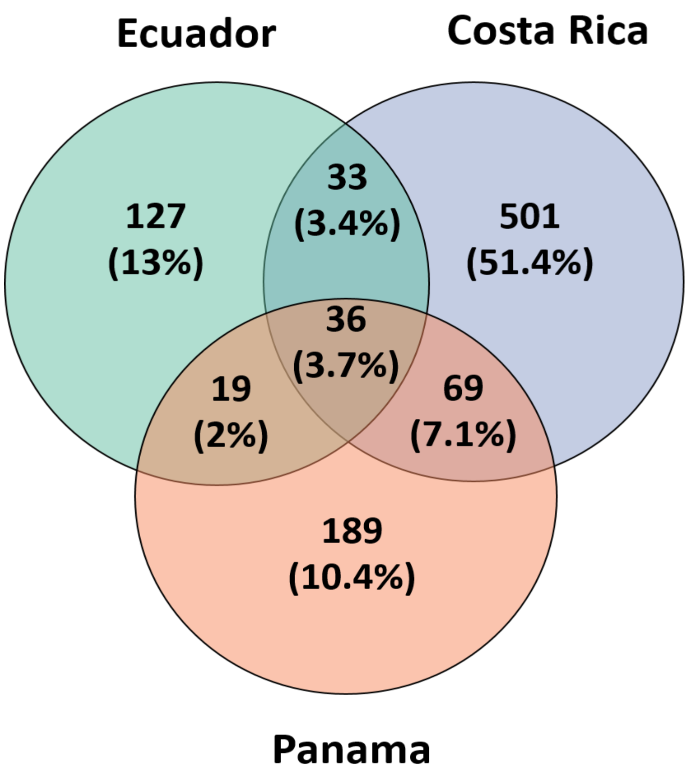

**Figure S2.** Number of Microsporidian OTUs exclusive and shared by country. Costa Rica has the highest number of exclusive OTUs. The number of shared Microsporidian OTUs between countries is very low. The Venn diagram was constructed using the online tool Venny 2.0 (Oliveros, 2007).
